# Notch Inhibition Enhances Morphological Reprogramming of microRNA-Induced Human Neurons

**DOI:** 10.1101/2024.01.12.575384

**Authors:** Kyle F. Burbach, Andrew S. Yoo

**Author notes:** **<u>Contact information</u>** Correspondence.

## Abstract

Although the importance of Notch signaling in brain development is well-known, its specific contribution to cellular reprogramming remains less defined. Here, we use microRNA-induced neurons that are directly reprogrammed from human fibroblasts to determine how Notch signaling contributes to neuronal identity. We found that inhibiting Notch signaling led to an increase in neurite extension, while activating Notch signaling had the opposite effect. Surprisingly, Notch inhibition during the first week of reprogramming was both necessary and sufficient to enhance neurite outgrowth at a later timepoint. This timeframe is when the reprogramming miRNAs, miR-9/9* and miR-124, primarily induce a post-mitotic state and erase fibroblast identity. Accordingly, transcriptomic analysis showed that the effect of Notch inhibition was likely due to improvements in fibroblast fate erasure and silencing of anti-neuronal genes. To this effect, we identify *MYLIP*, whose downregulation in response to Notch inhibition significantly promoted neurite outgrowth. Moreover, Notch inhibition resulted in cells with neuronal transcriptome signature defined by expressing long genes at a faster rate than the control, demonstrating the effect of accelerated fate erasure on neuronal fate acquisition. Our results demonstrate the critical role of Notch signaling in mediating morphological changes in miRNA-based neuronal reprogramming of human adult fibroblasts.

## Introduction

During neural development, neural progenitor cells activate Notch to keep themselves in a mitotic state (Lui et al., 2011). This behavior is important for creating the necessary number of neurons in the brain. One of the key regulators of Notch signaling in neurogenesis is *miR-9*, which is known to be pro-neuronal and a negative regulator of Notch (Roese-Koerner et al., 2017; Yoo et al., 2011). MiR-9 expression appears to be mutually exclusive with Notch activity, with mature neurons activating miR-9 while mature glia have active Notch (Lui *et al*., 2011; Patten et al., 2006). Radial glia, the neural progenitors during development, oscillate between the two states as they divide (Roese-Koerner *et al*., 2017; Stappert et al., 2015). Previous studies have leveraged this rationale in direct reprogramming protocols to generate neurons by blocking Notch cleavage with DAPT, a γ-secretase inhibitor (Zhang et al., 2015). Conversely, other studies applied gain-of-function Notch signaling by overexpressing the Notch Intracellular Domain (NICD) to generate neural stem cells (Cassady et al., 2014; Karow et al., 2018) or inhibit neurogenesis *in vivo* (Mall et al., 2017). Additionally, it has been shown that inhibiting Notch signaling with DAPT interferes with successful neural stem cell reprogramming (Xiao et al., 2018). Despite these findings, it remains necessary to understand how Notch signaling affects other neuronal reprogramming methods.

Here, we implemented neuronal conversion of human fibroblasts using the neurogenic activity of microRNAs, miR-9/9* and miR-124 (miR-9/9*-124) that can erase the fibroblast identity and directly induce a neuronal fate when ectopically expressed (Abernathy et al., 2017; Cates et al., 2021; Yoo et al., 2011). These microRNA-induced neurons (miNs) bypass stem cell intermediates, which allows the reprogramming cells to retain the epigenetic age signature of pre-reprogrammed cells (Abernathy et al., 2017; Huh et al., 2016; Victor et al., 2018). Previous work has shown that this reprogramming occurs in two distinct phases where miR-9/9*-124 first erase the fibroblast identity, followed by the acquisition of neuronal fate (Cates et al., 2021). Despite the importance of Notch signaling to neuronal fate determination *in vivo*, the role of Notch in direct neuronal reprogramming remains elusive. We therefore sought to investigate the effect of inhibiting Notch signaling during miRNA-mediated neuronal conversion of human fibroblasts.

## Results

### Inhibiting Notch signaling increases neurite outgrowth in miNs

Three independent human primary fibroblast lines (GM02171, AG13396, AG08379) were reprogrammed into neurons using the established protocol based on miR-9/9*-124 (Church et al., 2021). To assay changes in reprogramming efficiency more sensitively, we assessed cells at post-induction day (PID) 21, after they have acquired neuronal identity but before they are fully mature (Cates et al., 2021) (Figure 1B). To test the effect of Notch inhibition, we first treated reprogramming cells with the γ-secretase inhibitor DAPT (3uM) for the duration of reprogramming and compared the morphological features compared to DMSO (0.1%) control treatment. At PID 21, we observed a dramatic increase in neurite length, showing a 23-44% increase per cell across cell lines (Figure 1C). Because GM02171 showed the largest change, we focus on this line as the model for subsequent experiments.

**Figure 1.**
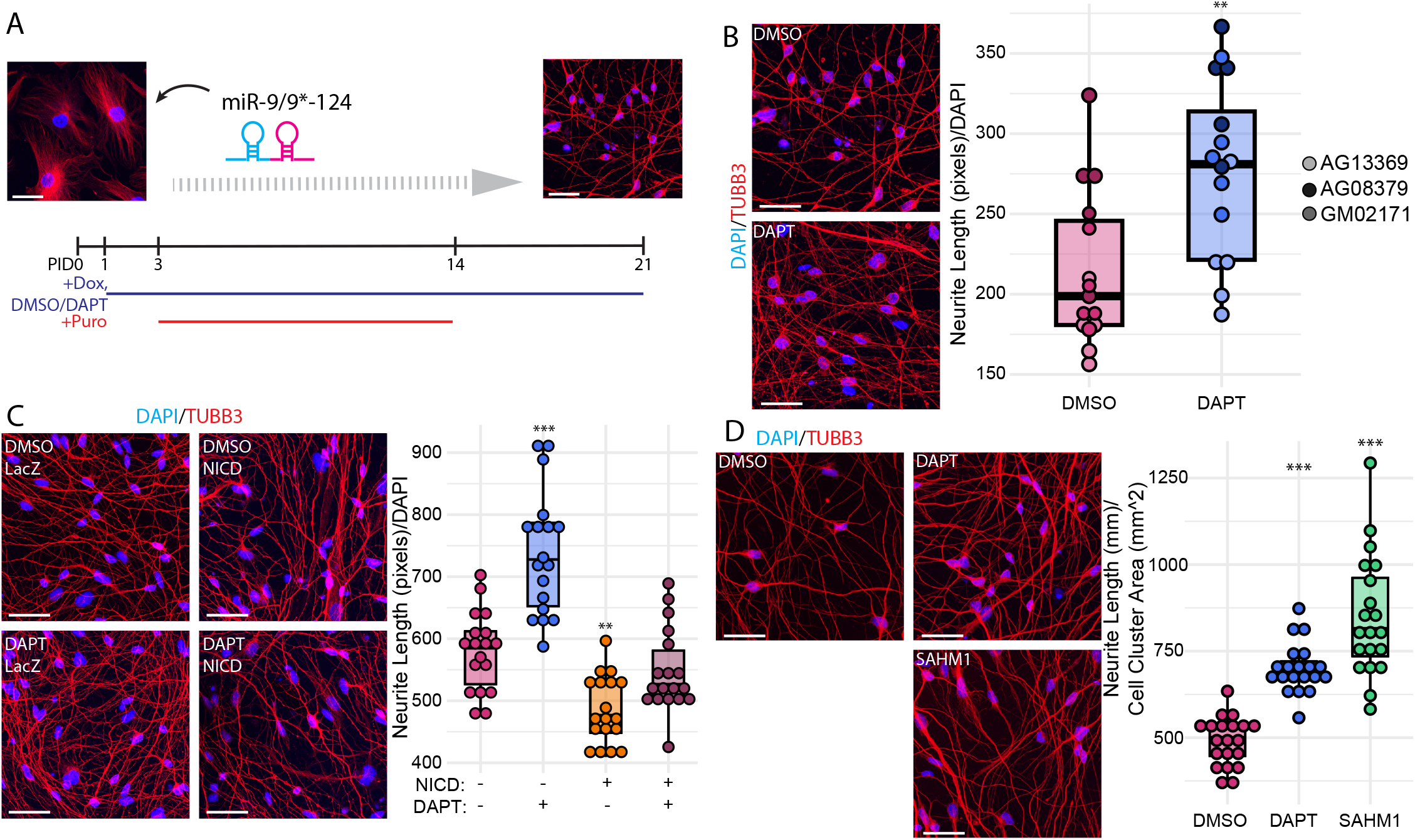
Inhibiting Notch signaling increases neurite outgrowth in miNs. A: Schematic of experimental design. After adding miR-9-124, cells are exposed to either DMSO or DAPT for 21 days, with Puro selection from days 3-14. B: Images showing beta-tubulin staining of fibroblasts, DMSO treated miNs, and DAPT treated miNs and quantification of neurite length per cell, n=3 cell lines, n=5 technical replicates (wells). C: Beta-tubulin staining of miNs treated with either DMSO or DAPT and overexpressing either LacZ or NICD and quantification of neurite length per cell, n=18 images across 3 wells. D: Beta-tubulin staining of miNs treated with DMSO, DAPT, or SAHM1 and quantification of neurite length per cell, n=20 images across 10 wells. Scale bars=50um.

DAPT inhibits Notch signaling by inhibiting γ-secretase, which in turn prevents the cleavage of Notch proteins and release of the Notch intracellular domain (NICD). However, there are at least 80 proteins that can be targeted by γ-secretase, including APP, DELTA1, JAGGED2, and ROBO1 (Haapasalo and Kovacs, 2011), suggesting that the observed phenotype could be due to reduced cleavage of any of the targets of γ-secretase. To confirm the specificity of Notch inhibition in enhancing neurite outgrowth of reprogrammed neurons, we overexpressed NICD and compared the extent of neurite extension to LacZ control overexpression. We found that NICD overexpression drastically reduced neurite length per cell compared to the LacZ control and that NICD in the presence of DAPT dampened the effect of DAPT (Figure 1D).

To further confirm the neurite phenotype is due specifically to Notch inhibition, we used the Notch Ternary Complex (NTC) inhibitor, SAHM1 (Moellering et al., 2009). The NTC consists of the NICD, RBPJ bound to the DNA, and MAML which recruits chromatin remodeling proteins (Fortini and Artavanis-Tsakonas, 1994; Petcherski and Kimble, 2000; Wu et al., 2000). SAHM1 is a small peptide which mimics an alpha helix in MAML, allowing it to exclude MAML from binding its groove in RBPJ (Moellering *et al*., 2009). This exclusion prevents the recruitment of chromatin remodelers and therefore inhibits the Notch transcriptional program. Addition of SAHM1 (2uM) resulted in miNs with a similar phenotype to DAPT-treated cells. When quantified through phase contrast and immunofluorescence, SAHM1 treatment resulted in even longer neurites per cell compared to DAPT (Figure 1E). Therefore, the finding that a specific NTC inhibitor SAHM1 phenocopied the effect of DAPT and NICD inhibited the neurite inhibition, demonstrates that the morphological enhancement of neurite extension by DAPT during miRNA-mediated reprogramming is due to Notch inhibition.

### Increased neurite length is due to transcriptional changes in the first week of reprogramming

Previous work has shown that microRNA-directed reprogramming occurs in two discrete steps: erasure of the original cell’s identity and acquisition of neuronal identity (Cates et al., 2021). To further investigate the mechanism behind how Notch inhibition results in improved neurite extension, DAPT was added for one-week intervals from PID0 to PID28 (1-7, 7-14, 14-21, 21-28), from PID0-21 as positive control, and DMSO as vehicle control. Live phase contrast imaging was performed using the Incucyte platform and neurite length was quantified using the Neurotrack module (Sartorius, 2016). Comparing between different treatment intervals, we found that DAPT had the strongest effect during the first seven days of reprogramming when the neurite length was assessed at PID28 (Figure 2B), similar to cells treated with DAPT for 21 days. These results indicate that Notch inhibition helps with the early reprogramming phase when miR-9/9*-124 erase the fibroblast identity.

**Figure 2.**
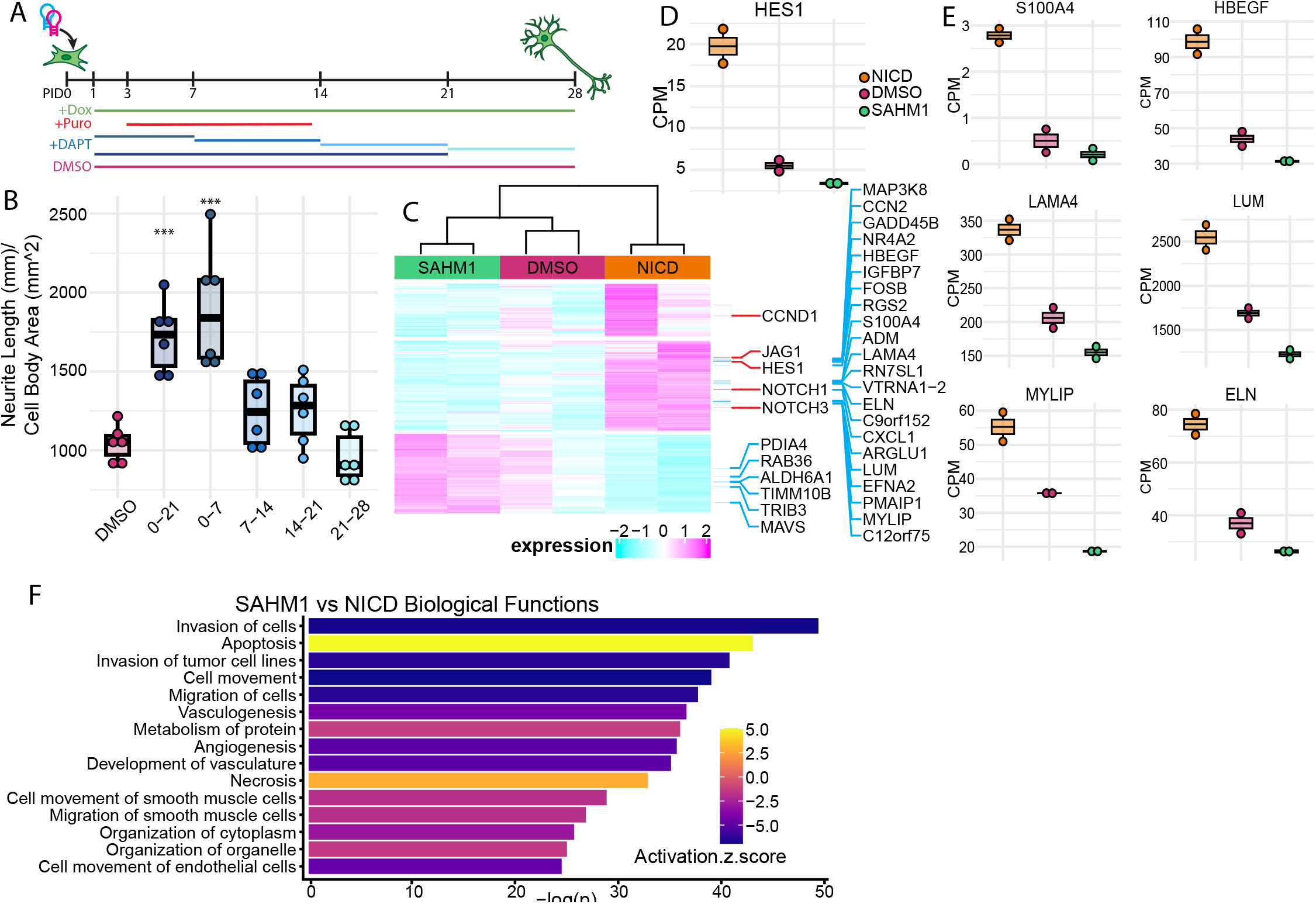
Increased neurite length is due to transcriptional changes in the first week of reprogramming. A: Schematic showing the experimental design. miNs were treated with DAPT for one-week intervals for four weeks. B: Quantification of neurite length for each condition, n=6 wells. C: Heatmap showing transcriptional changes because of changes to Notch signaling. Samples are sorted by Pearson correlation, and genes were grouped into seven clusters by k-means clustering. Shown clusters represent groups which are downregulated comparing SAHM1 to DMSO and upregulated comparing NICD to DMSO, or vice versa. Expression changes are Log2 fold change. D: IGV tracks showing changes in HES1 expression because of changing Notch activity. E: Box charts showing CPM for representative genes from cluster showing increased expression with Notch activation and decreased expression with Notch inhibition. Genes are either fibroblast expressed or anti-neuronal. F: Top 15 differential biological functions between SAHM1 and NICD analyzed by IPA.

To further investigate how Notch inhibition alters the early phase of reprogramming, we performed RNA-seq in reprogramming cells treated with DMSO or SAHM1 and overexpressing LacZ or NICD at PID6 (Referred to as DMSO, SAHM1, or NICD-treated). Differentially expressed genes (DEGs) were generated using DESeq2 (adj p-value≤0.05, |log2fold change|≥0.5). K-means clustering showed that SAHM1 and DMSO-treated cells were more similar than NICD-treated miNs (Figure 2C). Notch signaling genes and their targets show downregulation due to SAHM1 and upregulation due to NICD, as illustrated by HES1 (Figure 2D). A subset of DEGs which showed strong downregulation with Notch inhibition (SAHM1) and upregulation with Notch activation (NICD) or vice versa, were further examined to focus on genes that responded to differential Notch activities. This gave a list of 28 genes, which showed enrichment for fibroblast identity genes such as *S100A4, LAMA4, LUM, ELN*, and *HBEGF* (Figure 2E) and genes enriched in neural progenitors such as *FOSB, NR4A2, EFNA2, LINC01089* (Supplemental Figure 1A). This indicates that NICD interferes with miR-9/9*-124’s ability to erase the fibroblast identity and correctly establish neuronal identity.

Furthermore, we used Ingenuity Pathway Analysis (IPA) to analyze differentially affected biological functions. The top 15 most significant terms comparing SAHM1 to NICD show that aside from the expected Organization of Cytoplasm and Organization of Organelle, there is little to no consistency in the terms returned, and they span functions across a variety of non-neuronal tissues (Figure 2F). This is consistent with the notion that in the early phase of reprogramming, cells are primarily losing their original cell identity rather than simultaneously acquiring the neuronal fate – reflected by the lack of neuronal terms even within the top 50 terms (Supplemental Figure 1B). Therefore, our finding suggests that Notch inhibition specifically facilitates silencing of non-neuronal genes before reprogramming cells transition into the neuronal fate.

### MYLIP is a Notch target gene responsible for regulating neurite outgrowth

A particularly interesting DEG is *MYLIP*, which has been demonstrated to interfere with neural cells’ ability to grow neurites (Bornhauser et al., 2003; Olsson et al., 1999). Furthermore, ChIP of RBPJ binding sites identified a pair of binding sites upstream of and at the TSS of MYLIP, indicating it is a regulatory target of Notch (Supplemental Figure 1C) (Wang et al., 2011). Targetscan also identified a miR-124 binding site in the 3’UTR, indicating that it should be targeted by the microRNAs. However, our previously published time course RNA-seq (Abernathy *et al*., 2017) shows that MYLIP expression empirically increases starting at PID6 (Supplemental Figure 1C). To test the role of *MYLIP* in miNs, we knocked down *MYLIP* using shRNAs (Figure 3B). Even with the mild knockdown level of 25% reduction in *MYLIP* transcript, *MYLIP* shRNA resulted in a 43% increase in neurite length at PID21 (Figure 3C). Therefore, our results reveal *MYLIP* as a Notch-responsive gene whose repression is critical for neurite outgrowth of miNs.

**Figure 3.**
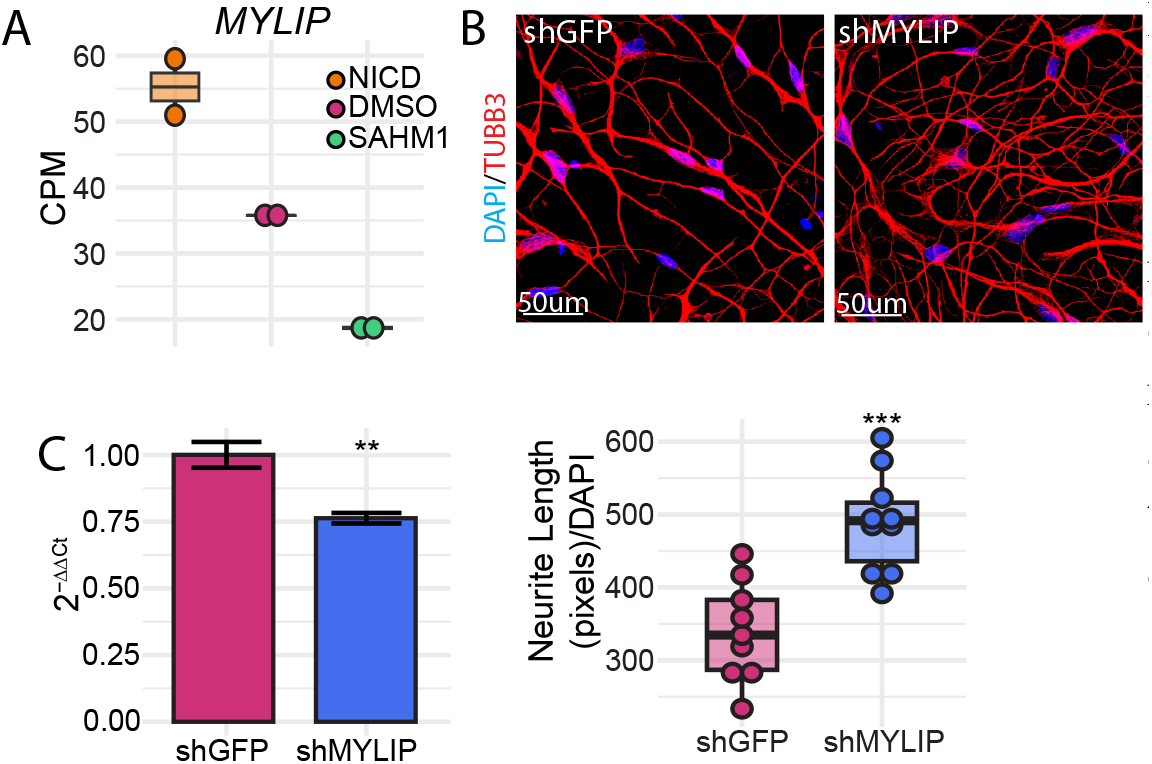
MYLIP is a Notch target gene responsible for regulating neurite outgrowth. A: Boxplot showing CPM for MYLIP in Notch overexpressing, DMSO treated, and SAHM1 treated miNs at PID7. Dots represent RNAseq replicates. B: Representative images showing miNs expressing shRNA against GFP or MYLIP. C: Left: qPCR results of MYLIP knockdown as 2-ΔΔCt, normalizing against GAPDH in shGFP treated miNs. Right: Quantification of neurite length per cell from B. Each dot represents one image taken from three coverslips.

### Long gene expression analysis demonstrates Notch Inhibition enhances neuronal conversion

To investigate the effects of Notch inhibition on miNs as they acquire neuronal identity, miNs treated with either DMSO, DAPT, or SAHM1 were collected at PID21 for RNA-seq. By looking for changes found in both the DAPT and SAHM1 conditions but not the DMSO condition, we searched for Notch specific effects contributing to neurite growth. This showed that most of the variance found in the dataset can be attributed to Notch inhibition, as PC1 and 2 account for over 98% of the variance and splits DAPT and SAHM1 from DMSO (Supplemental Figure 1D). We investigated transcripts at this timepoint by expression of genes by length, as well as differentially expressed genes (Figure 4A). Analysis of the expression of long genes, a transcriptomic feature unique in neurons (Gabel et al., 2015; McCoy et al., 2018), shows that miNs treated with Notch inhibitors have higher long gene expression compared to controls at PID21. This indicates that Notch inhibition is facilitating miNs to mature into neurons at a faster rate than the control. The heatmap shows that most differentially expressed long genes (LDEGs) fall into the upregulated with Notch inhibitors group (178 upregulated vs 83 downregulated). Looking at LDEGs individually, many neuronal genes, such as *CDH13, KCNMA1*, and *DCC* are upregulated by Notch inhibitors (Figure 4B).

**Figure 4.**
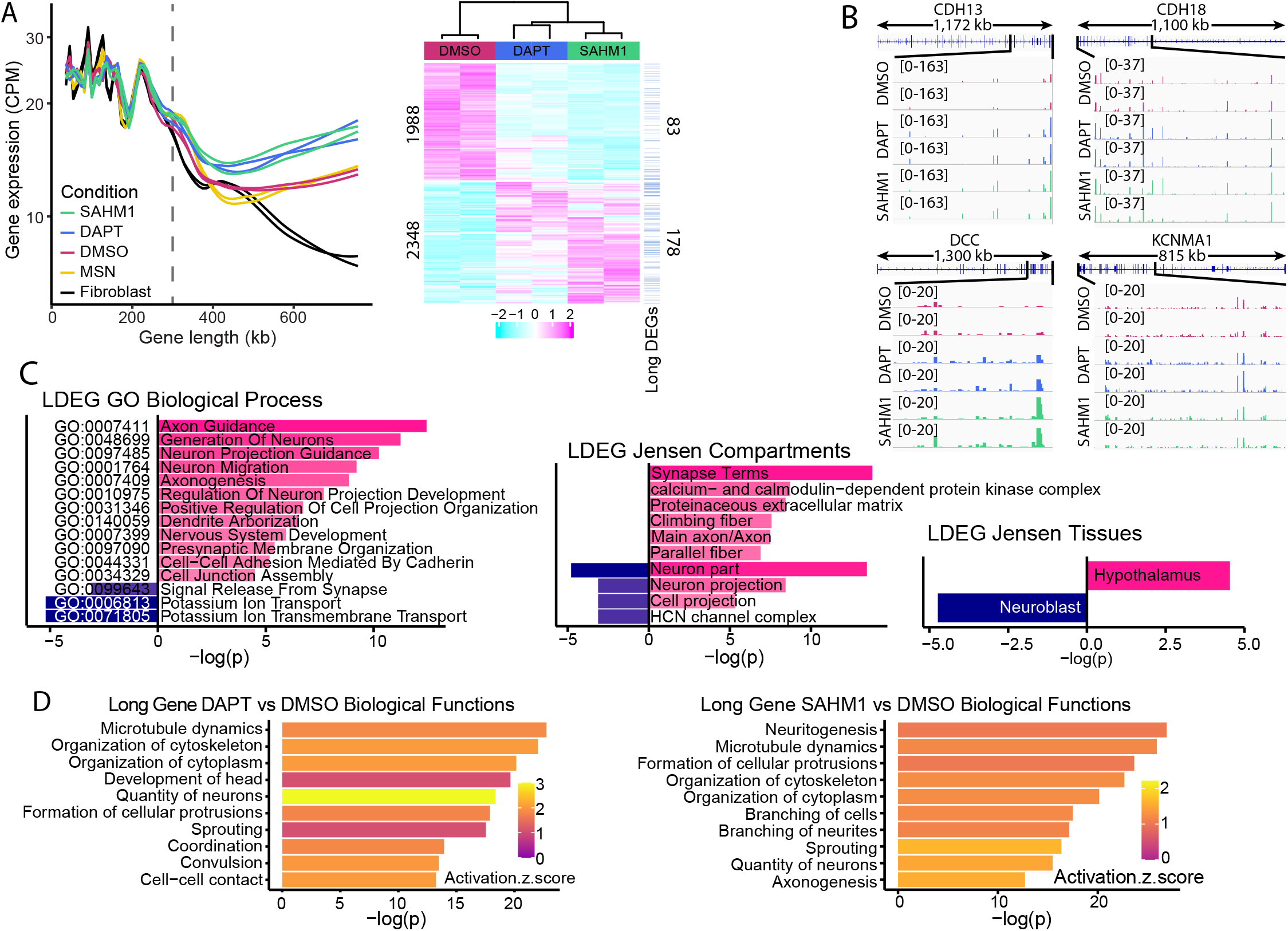
Long gene expression demonstrates Notch Inhibition improves miN reprogramming. A: Left; LONGO analysis on fibroblasts and miNs treated with DMSO, DAPT, or SAHM1. Right; Heatmap of RNAseq data. Samples are sorted by Pearson correlation, and genes were grouped into two clusters by k-means clustering. Expression changes are Log2 fold change. B: IGV tracks of representative neuronal long genes. C: ENRICHR analysis of LDEGs. Left; Go biological processes. Middle; Jensen Cellular Compartments. Right; Jensen Tissues. D: IPA differential biological functions analysis on LDEGs. Left; DAPT compared against DMSO. Right; SAHM1 compared against DMSO.

To further investigate the functions of these LDEGs, we ran the lists of 178 upregulated and 83 downregulated through Enrichr to determine their functions and importance to reprogramming (Chen et al., 2013; Kuleshov et al., 2016; Xie et al., 2021). Enrichr results were filtered for adjusted p-value ≤0.05 (Figure 4C). The top GO Biological Processes were filtered and shows that the upregulated LDEGs enrich for exclusively neuronal functions, particularly development and maturation of neurons. Downregulated LDEGs are also related to neuronal function, but specifically potassium transport and chemical signaling, indicating that this group of long genes are enriched for potassium and other ion channels. However, the downregulated ion channels appear to be consistently lowly expressed (CPM≤ 10), which likely suggests that they may not be critical for neuronal reprogramming. Jensen compartments shows that upregulated LDEGs are enriched for neuronal growth and synaptic function, while downregulated are mostly in neuron part and HCN channel complex, which as a voltage regulated potassium channel reinforces what we found with GO biological processes. Jensen tissues shows that the upregulated LDEGs identify only with the hypothalamus, a part of the mature brain, while downregulated LDEGs are only associated with neuroblasts, immature neurons still undergoing development. This further confirms that inhibiting Notch signaling improves the outcome of reprogramming miNs.

We performed additional pathway analysis using IPA on LDEGs. Comparing each inhibitor to DMSO, changes to neurite growth and neuronal function were confirmed by the presence of terms such neuritogenesis, microtubule dynamics, quantity of neurons, differentiation of neurons, branching of cells and formation of cellular protrusions (Figure 4D). This shows that although at PID7 cells were not specifically acquiring neuronal fate following Notch inhibition, at PID21 the effects which were enhancing fibroblast fate erasure are resulting in cells which are transcriptionally more neuronal than the control. These data indicate that inhibiting Notch in miN reprogramming results in neurons which express a more mature neuronal transcriptome indicated by neuronal long gene expression.

## Discussion

Using DAPT to inhibit Notch signaling in miNs provides an easy way to robustly improve neurite extension in reprogrammed neurons. Additionally, the Notch specific inhibitor SAHM1 allows for the same phenotype, but without the possibility of off targets that come with inhibiting γ-secretase. Notably, this is an important consideration in the study of neurodegenerative disorders such as Alzheimer’s Disease (AD). One of the key hallmarks of AD is accumulation of amyloid-β (Aβ) plaques. Aβ is one of the targets of γ-secretase, and therefore using DAPT would be inappropriate to study AD in reprogrammed neurons. However, being able to use SAHM1 or only temporarily inhibit Notch during early phase of reprogramming would enhance neuronal reprogramming without affecting APP processing. Meanwhile, as many of the phenotypes associated with AD are localized to the axons, such as tau tangles and dystrophic neurites, generating miNs with more robust neurite extension with SAHM1 application may enhance the manifestation of AD-associated phenotypes.

It is intriguing that addition of Notch inhibitor for the first week of reprogramming was sufficient to have a profound impact on the reprogramming outcome weeks later. Specifically, there is no robust transcriptional signature indicative of developing neurons at PID7, yet the early non-neuronal changes by Notch inhibition cascade into a strong, proneural transcriptome two weeks later. This finding reinforces the importance of effective erasure of the original cell’s identity leading to an adoption of neuronal identity, as also exemplified by MYT1L as a reprogramming factor (Mall *et al*., 2017). Cellular reprogramming is a complicated process involving many transcriptional and chromatin switches, and the ability for one switch early in reprogramming to have a dramatic effect on the final output highlights the importance of dissecting mechanisms underlying fate erasure during neuronal conversion.

By investigating the transcriptomic changes in reprogramming miNs in early and late phases with Notch inhibition, we have identified *MYLIP* as a gene whose repression aids in facilitating morphological changes in the form of neurite outgrowth. In neural development, *MYLIP* had been demonstrated to be important in the development of neurites in cells differentiating from stem cells (Bornhauser *et al*., 2003; Olsson *et al*., 1999). Demonstrating the importance of this same gene in direct neuronal conversion underscores some of the genetic pathways common to neurogenesis and cellular reprogramming. Therefore, future studies stemming from this study should aim to uncover other Notch-responsive genes with undefined roles in neurogenesis and neural development.

## Experimental Procedures

### MicroRNA-mediated neuronal reprogramming

The construction of all plasmids used in this study was previously described, and they are publicly available at Addgene as pTight-9-124-BclxL (60857), rtTA-N144 (66810), LacZ (42560), and TetO-FUW-NICD (61540) (Cassady et al., 2014; Church et al., 2021). shRNAs were bought from Sigma’s Mission Library (TRCN0000263127 and TRCN0000282500). Lenti-X 293T cells were acquired from Clontech (632180). Adult dermal fibroblasts were acquired from Coriell (GM02171, AG08379, AG11369).

Lentivirus was produced from each individual plasmid and pooled for transduction as previously described (Church et al., 2021; Richner et al., 2015). Lenti-X 293Ts were transduced with psPAX2, pMD2.G, and the corresponding packaging plasmid and polyethyleneimine (Polysciences). After 72 hours, supernatant was collected, filtered through 0.45-um PES membranes, and virus was concentrated by spinning at 70,000g for 2 hours at 4C. Lentivirus was resuspended in PBS and stored at -80C.

To reprogram fibroblasts into miNs, the rTTA and pTight-9-124-BclxL lentiviruses were added to fibroblasts with polybrene (Sigma-Aldrich, H9268) for 22-24 hours as previously published (Church et al., 2021). For relevant experiments, LacZ or TetO-FUW-NICD were also added. Cells were then washed and fresh 15% DMEM with 1ug/ml doxycycline (DOX) was added with the appropriate compound: DMSO (Millipore Sigma, D2650), DAPT (Millipore Sigma, D5942), or SAHM1 (Millipore Sigma, 491002). After two days the media was changed with the addition of 3mg/ml puromycin, then replated onto coverslips coated with fibronectin and laminin after another two days. On day six DMEM was removed and replaced with Neuronal Medium (NM, ScienCell, 1521) supplemented with 1mM valproic acid, 200mM dibutyl cAMP, 10ng/ml BDNF, 10ng/ml NT-3, 1mM RA, 200uM ascorbic acid, and 1x RevitaCell Supplement (RVC). Cells were half-fed this same medium every four days, with additional dox supplementation every other day in between half-feedings. RVC and puromycin were no longer added after day 14.

### Immunostaining

When ready, cells are fixed in 4% paraformaldehyde (Electron Microscopy Sciences, 15710) for 20 min. After washing three times with PBS, primary antibodies are added in a solution of 5% bovine serum albumin and 0.2% Triton-X. This incubates at 4C overnight. Cells are then washed three times with the BSA-TritonX solution without any antibodies, then secondary antibody is added for two hours at room temperature. Cells are then washed with BSA-TritonX once and PBS once, then incubated in DAPI (Sigma-Aldrich, D-9542) solution for 15 minutes before being placed back in PBS. A Leica SP5X white light laser confocal system with Leica Application Suite Advanced Fluorescence (LAS AF) v2.7.3.9723 was used to capture images.

Primary antibodies used include mouse anti-BETA TUBULIN III (Biolegend 801202), rabbit anti-BETA TUBULIN III (Biolegend 802001), and rabbit anti-MAP2 (Cell Signaling #4542). Secondary antibodies used include Alexa Fluor 488 goat anti-mouse IgG (H+L) (Invitrogen, A-11029, 1:2,000), Alexa Fluor 488 goat anti-rabbit IgG (H+L) (Invitrogen, A-11034, 1:2,000), Alexa Fluor 568 goat anti-mouse IgG (H+L) (Invitrogen, A-11031, 1:2,000), Alexa Fluor 568 goat anti-rabbit IgG (H+L) (Invitrogen, A-11016, 1:2,000), Alexa Fluor 594 goat anti-mouse IgG (H+L) (Invitrogen, A-11032, 1:2,000), Alexa Fluor 594 goat anti-rabbit IgG (H+L) (Invitrogen, A-11012, 1:2,000), and Alexa Fluor 594 goat anti-rat IgG (H+L) (Invitrogen, A-11007, 1:2,000).

### Quantification methods

Two methods were used to quantify neurite length. Confocal images were analyzed using Amira software. Live cell tracking was also used with the Incucyte S3 from Sartorius along with the Neurotrack module. Results were plotted using ggplot2 in R.

### RNA-seq and differential gene expression analysis

When ready, RNA was extracted from miNs using the RNAeasy Micro/Mini Kit (Qiagen). Purified RNA was submitted to the Genome Access Technology Center at Washington University to prepare libraries and sequence. Samples were prepared according to library kit manufacturer’s protocol, indexed, pooled, and sequenced on an Illumina NovoSeq. Basecalls and demultiplexing were performed with Illumina’s bcl2fastq software and a custom python demultiplexing program with a maximum of one mismatch in the indexing read. RNA-seq reads were then aligned to the Ensembl release 76 primary assembly with STAR version 2.5.1a (Dobin et al., 2013). Gene counts were derived from the number of uniquely aligned unambiguous reads by Subread:featureCount version 1.4.6-p5 (Liao et al., 2014). Isoform expression of known Ensembl transcripts were estimated with Salmon version 0.8.2 (Patro et al., 2017). Sequencing performance was assessed for the total number of aligned reads, total number of uniquely aligned reads, and features detected. The ribosomal fraction, known junction saturation, and read distribution over known gene models were quantified with RSeQC version 2.6.2 (Wang et al., 2012).

All gene counts were then imported into the R/Bioconductor package EdgeR and TMM normalization size factors were calculated to adjust for samples for differences in library size (Robinson et al., 2010). Ribosomal genes and genes not expressed in the smallest group size minus one and samples greater than one count-per-million were excluded from further analysis. The TMM size factors and the matrix of counts were then imported into the R/Bioconductor package Limma (Ritchie et al., 2015). Weighted likelihoods based on the observed mean-variance relationship of every gene and sample were then calculated for all samples with the voomWithQualityWeights (Liu et al., 2015). The performance of all genes was assessed with plots of the residual standard deviation of every gene to their average log-count with a robustly fitted trend line of the residuals. Differential expression analysis was then performed to analyze for differences between conditions and the results were filtered for only those genes with Benjamini-Hochberg false-discovery rate adjusted p-values less than or equal to 0.05.

For each contrast extracted with Limma, global perturbations in known Gene Ontology (GO) terms, MSigDb, and KEGG pathways were detected using the R/Bioconductor package GAGE to test for changes in expression of the reported log 2 fold-changes reported by Limma in each term versus the background log 2 fold-changes of all genes found outside the respective term (Luo et al., 2009). The R/Bioconductor package heatmap3 was used to display heatmaps across groups of samples for each GO or MSigDb term with a Benjamini-Hochberg false-discovery rate adjusted p-value less than or equal to 0.05 (Zhao et al., 2014). Perturbed KEGG pathways where the observed log 2 fold-changes of genes within the term were significantly perturbed in a single-direction versus background or in any direction compared to other genes within a given term with p-values less than or equal to 0.05 were rendered as annotated KEGG graphs with the R/Bioconductor package Pathview (Luo and Brouwer, 2013).

To find the most critical genes, the raw counts were variance stabilized with the R/Bioconductor package DESeq2 and was then analyzed via weighted gene correlation network analysis with the R/Bioconductor package WGCNA (Langfelder and Horvath, 2008; Love et al., 2014). Briefly, all genes were correlated across each other by Pearson correlations and clustered by expression similarity into unsigned modules using a power threshold empirically determined from the data. An eigengene was then created for each de novo cluster and its expression profile was then correlated across all coefficients of the model matrix. Because these clusters of genes were created by expression profile rather than known functional similarity, the clustered modules were given the names of random colors where grey is the only module that has any pre-existing definition of containing genes that do not cluster well with others. These de-novo clustered genes were then tested for functional enrichment of known GO terms with hypergeometric tests available in the R/Bioconductor package clusterProfiler (Yu et al., 2012). Significant terms with Benjamini-Hochberg adjusted p-values less than 0.05 were then collapsed by similarity into clusterProfiler category network plots to display the most significant terms for each module of hub genes in order to interpolate the function of each significant module. The information for all clustered genes for each module were then combined with their respective statistical significance results from Limma to determine whether or not those features were also found to be significantly differentially expressed.

DEGs were run through IPA (Qiagen) Core Analysis with minimum thresholds of adjusted p-value≤0.05 and |log2fold change|≥0.5, increasing the stringency if more than 8000 genes survived filtering.

### Statistics and reproducibility

Staining and quantification were initially done with three cell lines were repeated five times. Subsequent experiments focused on one cell line with the largest effect, with at least three experiments. Statistical tests were performed in R as either a two-tailed unpaired t-test or one-way ANOVA with post hoc Tukey’s test. Data was tested for normality using the Shapiro test and for heterogeneity using the Levene test, and appropriate corrections were done for non-heterogeneous data. No data was excluded from analysis. Significance is defined as <=0.05, **<=0.01, ***<=0.01. Analyses were automated whenever possible to eliminate bias, and samples were blinded when manual counting was required. Two technical replicates were used for RNAseq analysis, which showed highly reproducible results. RNAseq analysis used a cutoff for adjusted p-values <= 0.05.

## Resource availability

### Corresponding authors

Further information and requests for resources and reagents should be directed to and will be fulfilled by the corresponding author, Andrew Yoo.

### Materials availability

This study did not generate new unique reagents.

### Data and code availability

Data from this project are available at Gene Expression Omnibus (GEO: GSE252626 & GSE252733).

## Acknowledgements

We thank the Genome Technology Access Center at the McDonnell Genome Institute at Washington University School of Medicine for help with genomic analysis. The Center is partially supported by NCI Cancer Center Support Grant #P30 CA91842 to the Siteman Cancer Center. This publication is solely the responsibility of the authors and does not necessarily represent the official view of NCRR or NIH.

## Author Contributions

K.F.B. conducted the experiments and analyzed the results, and K.F.B. and A.S.Y. designed the experiments and wrote the paper. A.S.Y. supervised the project.

---

**Supplementary Figure 1.**
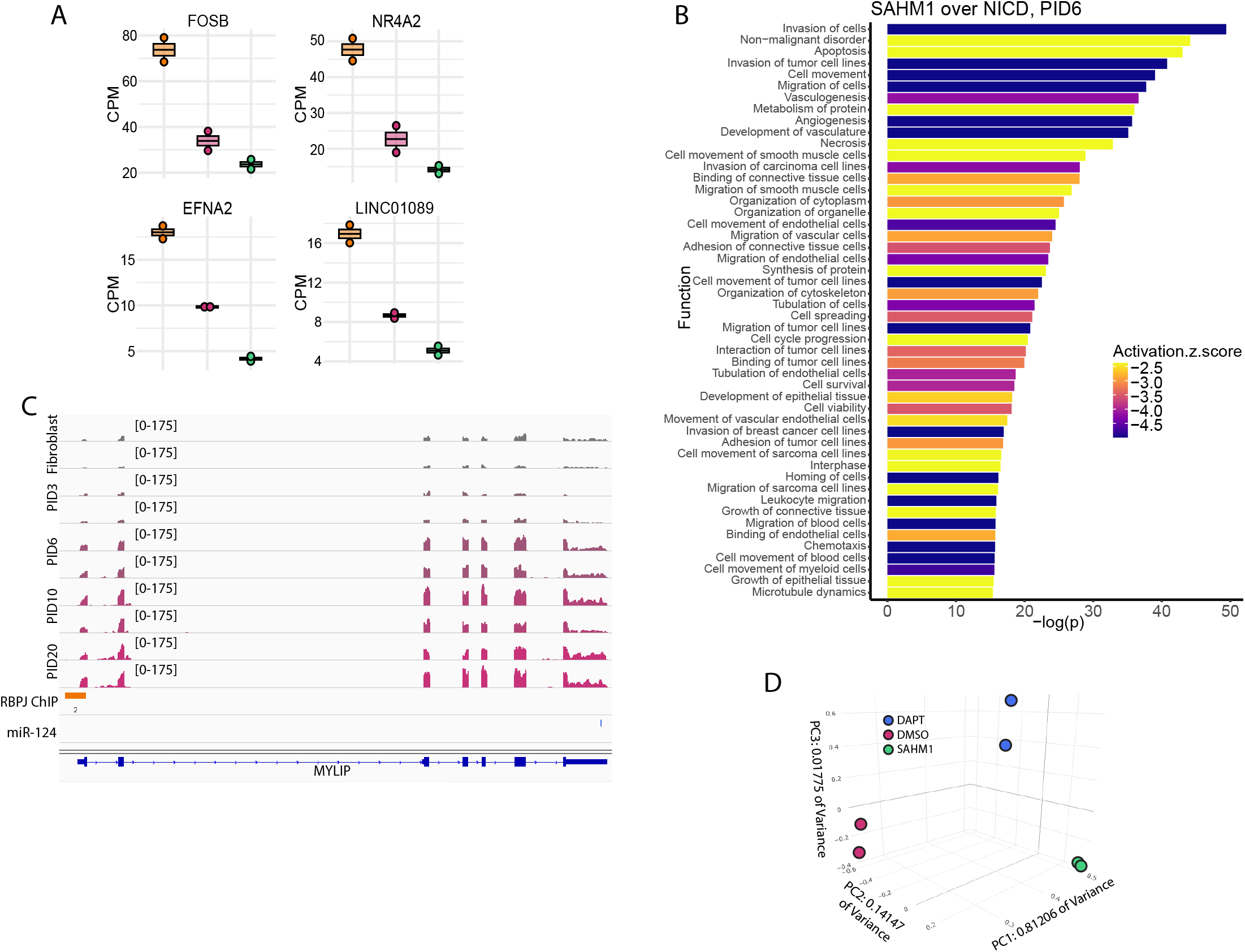
A: Bar charts showing CPM values for neuronal progenitor genes showing differential behavior because of Notch signaling in miNs at PID6 with NICD overexpression, SAHM1 treatment, or negative control. B: Top 50 biological function terms from PID6 comparing SAHM1 treated miNs to NICD overexpressing miNs, extended from figure 2F. C: IGV screenshot showing MYLIP expression at multiple timepoints during miN reprogramming, RBPJ ChIP data, and Targetscan predicted binding sites for miR-124. Targetscan did not predict any binding sites for miR-9. D: PCA showing first three principal components of miNs at PID21 treated with DMSO, DAPT, or SAHM1.

